# A Low-Cost Biological Agglutination Assay for Medical Diagnostic Applications

**DOI:** 10.1101/411637

**Authors:** Nicolas Kylilis, Pinpunya Riangrungroj, Hung-En Lai, Valencio Salema, Luis Ángel Fernández, Guy-Bart V Stan, Paul S Freemont, Karen M Polizzi

## Abstract

Affordable, easy-to-use diagnostic tests that can be readily deployed for point-of-care (POC) testing are key in addressing challenges in the diagnosis of medical conditions and for improving global health in general. Ideally, POC diagnostic tests should be highly selective for the biomarker, user-friendly, have a flexible design architecture and a low cost of production. Here we developed a novel agglutination assay based on whole *E. coli* cells surface-displaying nanobodies which bind selectively to a target protein analyte. As a proof-of-concept, we show the feasibility of this design as a new diagnostic platform by the detection of a model analyte at nanomolar concentrations. Moreover, we show that the design architecture is flexible by building assays optimized to detect a range of model analyte concentrations supported using straight-forward design rules and a mathematical model. Finally, we re-engineer *E. coli* cells for the detection of a medically relevant biomarker by the display of two different antibodies against the human fibrinogen and demonstrate a detection limit as low as 10 pM in diluted human plasma. Overall, we demonstrate that our agglutination technology fulfills the requirement of POC testing by combining low-cost nanobody production, customizable detection range and low detection limits. This technology has the potential to produce affordable diagnostics for both field-testing in the developing world, emergency or disaster relief sites as well as routine medical testing and personalized medicine.

## INTRODUCTION

Affordable point-of-testing diagnostic technology applied to resource-limited sites is one of the most promising biotechnologies for improving global health (Daar et al. 2002). The development of new diagnostics and their successful adoption in the field by end users requires consideration of various diverse factors including, but not limited to, scientific challenges, economic restrictions and practical considerations (Giljohann and Mirkin 2009). High impact technology will allow sensitive and specific detection, it will be low cost and portable to ensure accessibility, and it will have a user-friendly design that does not rely on sophisticated equipment (Urdea et al. 2006).

Immunoassays are a dominant technology in *in vitro* diagnostics (Borrebaeck 2000). Antibodies can be generated for the target analyte with very high specificity and relative ease. Among the various types of immunoassay formats for the development of rapid diagnostic tests, latex agglutination tests (LAT), which use antibody molecules immobilized on latex particles to detect the presence of an analyte, are available for more than 300 diseases and biomolecules (Ortega-Vinuesa and Bastos-Gonzalez 2001). Multivalent immuno-latex particles can recognize their target analyte molecules through specific interactions and form higher order complexes (Ortega-Vinuesa and Bastos-Gonzalez 2001). As the agglutination reaction proceeds, extensive cross-linking between analyte molecules and latex particles leads to the formation of very large complexes that can be visibly detected by the naked eye or monitored spectrophotometrically (Price 2001).

The use of whole cells as a bioanalytical platform for *in vitro* medical diagnostics offers a number of favorable characteristics such as low cost of production. Whole cells are self-replicating and can manufacture recognition elements such as antibodies, eliminating the need for expensive purification steps. In addition, as living organisms, whole cells can provide physiologically relevant data on the bioavailability of the analyte (van der Meer and Belkin 2010). Bacteria, especially model organisms such as *Escherichia coli*, are often used as bioreporters as they are amenable to genetic engineering and easy to culture. Bacterial whole cell biosensors have been genetically engineered to detect medically relevant analytes such as metabolites (Goers et al. 2017), toxic chemicals (Horswell and Dickson 2003) and mercury in urine (Roda et al. 2001), hydroxylated polychlorinated biphenyls in serum (Turner et al. 2007) and nitrogen oxides in both serum and urine samples (Courbet et al. 2015). Whole-cell biosensors usually rely on intracellular detection of the target analyte (Goers et al. 2013), but such designs are limited to the detection of analytes that can diffuse or be actively transported across the bacterial cell membrane. Nevertheless, there is a multitude of medically relevant analytes such as protein biomarkers of disease state (Assicot et al. 1993; Haverkate et al. 1997; Ohman et al. 1996; Thompson et al. 2004) that cannot cross the membrane barrier. A few bacterial whole-cell bioreporter designs have been demonstrated that detect the analyte extracellularly (Looger et al. 2003; Webb et al. 2016), but these are of limited versatility in terms of choice of target because of their mechanism of action.

Merging bacterial surface display technology and antibody engineering can provide a mechanism for the detection of extracellular analytes with whole-cell biosensors in a biological equivalent of the latex agglutination test (LAT). Single chain antibody fragments (scFv) have been displayed on the surface of bacteria using engineered display proteins that localize to the outer membrane of the cell (Daugherty et al. 1998; Francisco et al. 1993; Fuchs et al. 1991). However, scFvs often have poor solubility, misfold and aggregate when expressed in bacterial hosts. As an alternative, single domain antibodies or nanobodies, are antibody fragments from heavy chain-only antibodies produced by the *Camelidae* family or cartilaginous fish. They have high stability when expressed in bacterial hosts (Muyldermans 2013; Salema and Fernandez 2017) and have been demonstrated to functionally display on the outer membrane of *E. coli* (Salema et al. 2016a; Salema et al. 2016b; Salema et al. 2013) and other bacterial species (Cavallari 2017; Fleetwood et al. 2013).

Here we demonstrate the development of a biological equivalent of the LAT that utilizes the *E. coli* surface display of camelid nanobodies as a detection element. As a starting point, we used a nanobody against green fluorescent protein (GFP) with a tandem dimeric GFP (tdGFP) analyte to induce cross-linking as a proof-of-concept. Using this convenient platform and a mathematical model of the agglutination process, we explored the effects of variables such as the number of bacterial particles in the assay and the nanobody expression level on the limit of detection of the assay. The total number of nanobodies in the reaction was found to be the primary factor influencing the limit of detection. With this understanding, we then explored the modularity of the platform by creating a whole-cell biosensor for the detection of human fibrinogen (hFib), a clinically relevant biomarker high levels of which are associated with cardiovascular disease and inflammation (Willis et al. 2004) and low levels of which can be used in the diagnosis of disseminated blood clotting (Lowe et al. 2004). Here, we propose a modular biosensing platform that harnesses the versatility and sensitivity of the LAT, but without the requirement for expensive steps to isolate and purify antibodies.

## MATERIALS AND METHODS

### Strains, plasmids and cell culture conditions

*Escherichia coli* DH10B^™^ (F^-^ *mcr*A δ(*mrr*-*hsd*RMS-*mcr*BC) f80*lac*ZδM15 δ*lac*X74 *rec*A1 *end*A1 *ara*D139 δ(*ara-leu*)7697 *gal*U *gal*K λ^-^ *rps*L(Str^R^) was used in this study. The pNVgfp (Salema et al. 2013) and pNVFIB1-2 plasmids (Salema et al. 2016a), which express the anti-GFP nanobody and two variants of an anti-human fibrinogen nanobody surface displayed through a β-intimin, respectively, are described elsewhere. The pNVgfp02 - 10 constructs were assembled by standard cloning procedures from the pNVgfp construct by replacing the T7 promoter of the original plasmid with a BBa_J2310x constitutive promoter and a designed RBS element.

For nanobody expression from pNVgfp and pNVFIB1-2, a single colony of freshly transformed DH10B cells was inoculated into LB medium supplemented with 34 μg/mL chloramphenicol and grown for 2 hours at 37°C with shaking at 250 rpm. For anti-GFP nanobody expression cell cultures were induced with 1mM IPTG and incubated at 30°C for 3 hours and 250 rpm shaking. Cells were kept overnight at 4°C, then harvested by centrifugation, washed twice and re-suspended in sterile phosphate buffered saline (PBS) pH 7.4 to a final OD600 of 1 (∼1*10^9^ cells). The pNVgfp02-10 library of variants utilizes constitutive promoters, and in this case, cell cultures were not induced with IPTG, but instead were grown for 5 hours at 37°C with shaking at 250 rpm. For anti-fibrinogen nanobody expression, cell cultures were induced with 1mM IPTG and incubated overnight at 37°C with shaking at 250 rpm before cells were harvested by centrifugation. Cells were washed twice and re-suspended in sterile phosphate buffered saline (PBS) pH 7.4 to a final OD600 of 1 (∼1*10^9^ cells).

### Analysis of nanobody surface expression

100 μL of the cell suspension was incubated with 200 pmol of purified GFPmut3b at room temperature for 30 min (for pNVgfp) or 2 μL of 50 μg/mL Myc-Tag (9B11) Mouse mAb (Alexa Fluor ^®^ 488 conjugate, Cell Signalling Technology, Danvers, MA) at 4°C in the dark for 30 min (for pNVFIB). Cells were washed twice and resuspended in 100 μl PBS. Subsequently, 2 μL of labeled cells were placed on a microscope slide and examined under a Nikon Eclipse Ti inverted microscope (Nikon Instruments, Japan) with exposure time at 100 ms using a magnification of 60x with a Ph1 filter.

### Nanobody quantification

The levels of cell-surface displayed nanobody were determined indirectly by measuring the cell fluorescence of bacterial cell populations labeled with different concentrations of purified GFPmut3b. 100 µL of the cell suspension was incubated with various amounts of purified GFP (0-40 pmol) at room temperature for 30 min. Subsequently, cells were washed twice and resuspended in 100 μL PBS. The labeled cells were diluted 1:100 in PBS and analyzed by flow cytometry (Attune NxT flow cytometer, Thermo Fisher Scientific, USA) with at least 100,000 events. Samples were measured at excitation 488 nm and emission 530/30 nm, and data analysis from three replicates was performed using FlowJo software (vX.0.7, FlowJo, LLC, USA). Recorded events were gated using the ellipse gate function to select for single cells at forward scatter (FSC-H) values 629 – 23,000 and side scatter (SSC-H) values 6,300 – 58,000. These values were chosen for providing a large coverage of the cell population events recorded. The standard deviation of the mean cell fluorescence for each sample was calculated using Prism 7 software tool (GraphPad, USA) using geometric mean fluorescence values from three technical replicates of the flow cytometry measurements.

### Biological LAT

200 µL of the cell suspension was incubated with 20 µL-50 µL of various concentrations of target molecules in a clear, round bottom 96-well plate (Costar ^®^, USA). Prepared microplates were statically incubated for ∼ 18 hours at room temperature. Assay results were documented using a photographic camera in one or more of the following photography setups: i) a microplate was placed on top of a dark surface, ii) a microplate was placed on top of a blue LED transilluminator without the use of blue light filter and iii) a microplate was placed on top of a blue LED transilluminator and the screen for blue light absorption was placed on top of the microplate. As for the results testing with hFib, a microplate was placed on top of a light box and photos were taken at three to six wells per frame using a mobile phone.

### Fibrinogen mock samples

Human fibrinogen (hFib) (Plasminogen, vWF, and Fibronectin depleted; FIB3) was purchased from Enzyme Research Laboratories (USA) and diluted in PBS or in fibrinogen-deficient human plasma (Sekisui Diagnostics GmbH, Germany). For the plasma-based assays, fibrinogen stock solutions were made at ∼ 5 mg/mL in plasma and then diluted to 1 mg/mL in PBS.

### Mathematical model and computer simulations

The derivation of the mathematical model is described in Supplementary Note 3. The model was implemented in Matlab ® software using a third-party script (Garcia-Molla 2017a, b) for system simulations and sensitivity analysis.

## RESULTS & DISCUSSION

### Proof-of-concept for a new diagnostic assay system

For the detection of protein analytes, an assay format was adopted that resembles the format of the LAT. The assay design utilizes bacterial cells displaying a nanobody on their cell surface to facilitate analyte recognition in the extracellular environment (Figure 1A). The display of several copies of the nanobody results in an engineered bacterial cell multivalent for its target analyte. In the presence of di- or multi-valent analyte molecules or when bacterial cells expressing nanobodies against multiple epitopes of a monovalent antigen are mixed, cross-linking between cells and analyte molecules produces aggregated bacterial clumps. This agglutination of bacterial cells results from the specific binding of protein analyte molecules to displayed nanobody elements in a bacterium-protein-bacterium sandwich. It was expected that the behavior of this biological LAT would resemble the general curve exhibited in agglutination immunoassays (Figure 1B) where maximum agglutination of bacterial cells would be observed at a specific ratio of cells to antigenic molecules with deviation from this ratio leading to progressively lower levels of agglutination. The visual output of a lack (or low levels) of bacterial agglutination is a dense bacterial pellet, while high levels of agglutination result in a network or membrane-like structure that spreads across the bottom of a well when the reaction is carried out in round bottom microplate well (Figure 1C).

**Figure 1:**
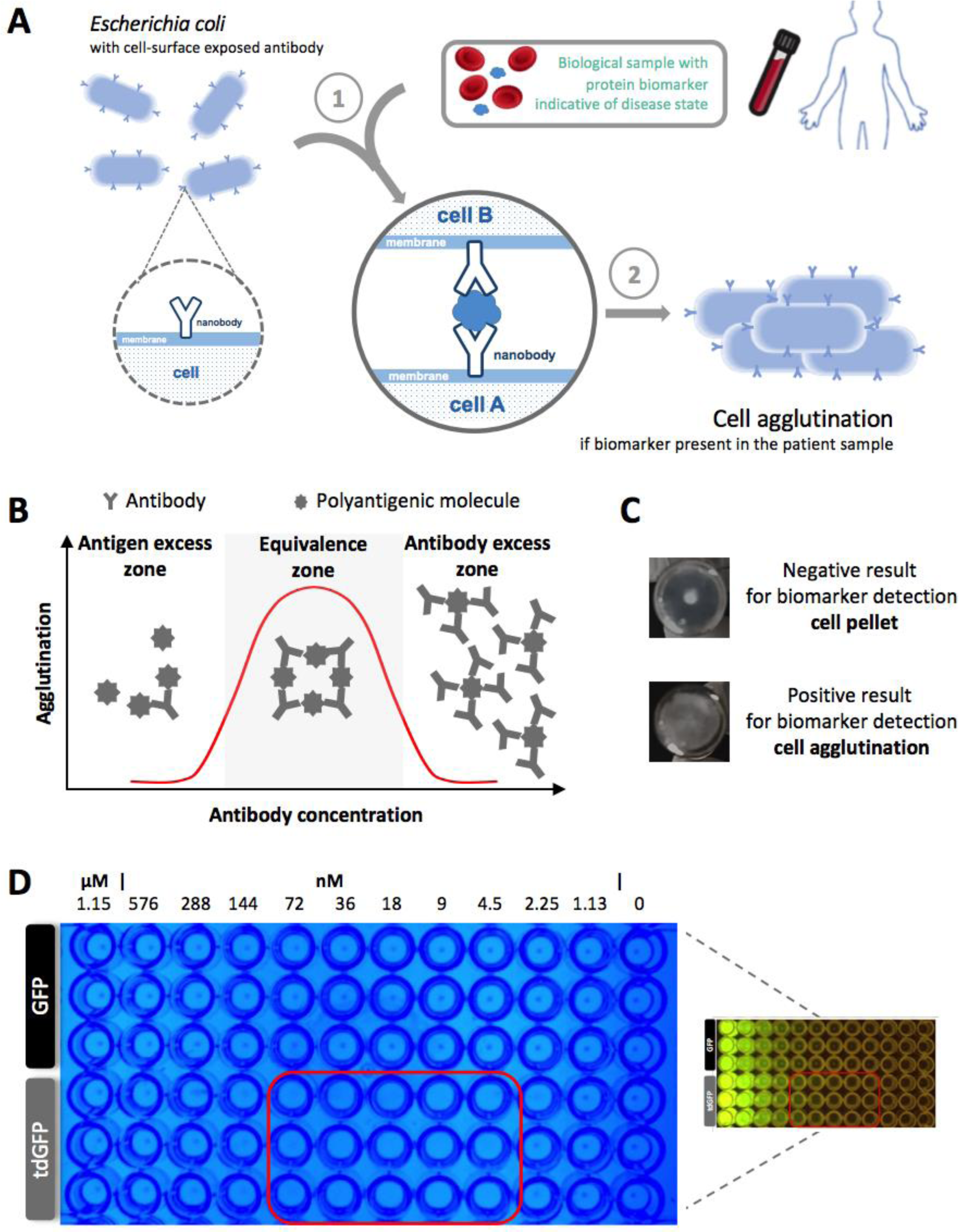
Low-cost, protein biomarker test for point-of-care medical diagnostics. **A** Assay design and test workflow for point-of-care testing. *E. coli* cells with surface-displayed nanobodies are mixed with a biological sample (e.g. blood) from a patient. If the patient sample includes the protein biomarker indicative of disease state, this will cause cross-linking between bacterial cells and biomarker molecules that leads to cell agglutination and changes the visual appearance of the test sample (test result). **B** The Heidelberger-Kendall curve describes the extent of aggregation of molecules in an agglutination reaction as a function of the antigen/antibodies ratio. **C** Positive and negative test results for cell agglutination assay. **D** Images of the proof-of-concept experiment, where *E. coli* cells displaying the anti-GFP nanobody on their cell surface have been incubated with monomeric GFP (negative control) or tdGFP (model biomarker) in microplates. The red box highlights wells with tdGFP concentrations that demonstrate high-levels of cell agglutination. In the main image, the microplate was back-illuminated with visible blue light for better contrast. In the secondary image, the same microplate was illuminated with the application of a GFP filter to demonstrate protein (GFP or tdGFP) concentrations.

For proof-of-concept experiments, *E. coli* cells were designed to express anti-GFP nanobody that is surface displayed through a β-intimin anchor (Salema et al. 2013). Surface localization and functional display of the anti-GFP nanobody was demonstrated by live cell fluorescence labeling and flow cytometry (Supplementary Note 1 and Supplementary Figure 1A-B) and the number of nanobodies displayed per cell was estimated by GFP titration followed by flow cytometry analysis (Supplementary Note 1 and Supplementary Figure 1C). To induce crosslinking of cells, a divalent analyte, tdGFP, was engineered by fusing a linker domain of 12 amino acids to two GFP monomers (Supplementary Note 2). Each domain of tdGFP can bind to an individual nanobody on separate cells initiating cell agglutination. To test this, we overexpressed and purified recombinant tdGFP (Supplementary Figure 2A). Fluorescence assays suggested that both GFP domains were correctly folded as the fluorescence intensity is approximately twice that of the GFP monomer at the same molarity (Supplementary Figure 2B).

To set up the agglutination assay, suspensions of *E. coli* cells expressing the anti-GFP nanobody were titrated with tdGFP analyte in concentrations ranging from 1.15 *μ*M to 1.13 nM in two-fold dilutions in round-bottom 96-well microplates. This experimental setup is typical of diagnostic hemagglutination assays taking place in clinical settings (Network 2011). After overnight incubation, cell agglutination was clearly visible in the analyte range of 4.5 nM to 75 nM tdGFP, in the form of a membrane-like structure across the bottom of the wells (Figure 1D). At lower (pre-zone) and higher (post-zone) tdGFP concentrations, distinct cell pellets were formed at the bottom of the wells (Figure 1D) indicating a lack of cell cross-linking. An analogous control reaction where cells were titrated with the equivalent concentrations of monomeric GFP, which should not initiate cell agglutination, showed cell pellet formation across the entire GFP analyte concentration range tested, indicating that cell agglutination is specific for the tdGFP analyte.

### Modulating the dynamic range of the diagnostic assay

The use of bacterial agglutination assay as a new diagnostic technology would benefit from designs that allow versatility in the detection of disease biomarkers. While the design of the demonstrated prototype assay platform is inherently modular since the nanobody can be engineered to bind specifically to any target, different medical conditions may require detection of molecular targets at different physiological concentrations in biological samples (Assicot et al. 1993; Haverkate et al. 1997; NICE 2014; Ohman et al. 1996; Thompson et al. 2004).

To enable further understanding of our nanobody-induced cell agglutination system and allow for the development of tunable assays for specific medical applications, we adapted a previously described mathematical model of the agglutination process (Dolgosheina et al. 1992) to the specifics of our technology. Briefly, our mathematical model describes the dynamics of cell agglutination in the presence of freely-diffusing analytes in which a finite number of cells are cross-linked via the tdGFP analyte under certain conditions, and consequently form larger aggregates over time. The output of the model is a measure of the expected level of bacterial cell agglutination at the end of the reaction compared to the initial number of free bacterial cells in the reaction, which we refer to as the normalized cell agglutination value 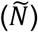 The model output is dependent on several variables which can be directly controlled in the laboratory such as the antibody binding rate constant (*C2*) and the dissociation rate constant (*C3*), the added concentration of analyte (*N*_l_), the number of nanobodies displayed per cell (*f*) and the concentration of cells in the reaction (*N*_0_). A more detailed explanation and derivation of the model, including a list of assumptions, can be found in Supplementary Note 3.

Initially, we tested the predictive capabilities of the model by simulating the cell agglutination reaction using variable values derived experimentally or from the literature (Supplementary Table 1). The simulation predicted that the maximum value of normalized cell agglutination would occur at 7 nM tdGFP analyte (Figure 2A, region between red dashed lines), in good agreement with the experimentally observed agglutination zone (Figure 2A, red rectangle). These results demonstrated that the model successfully captures the agglutination reaction dynamics. Next, we applied sensitivity analysis to the model to identify parameters with the largest influence on the agglutination reaction and the normalized cell agglutination value. The parameters that showed the greatest relative sensitivity were the normalized agglutination rate constant, *C*_1*n*_, and the total number of nanobodies in the reaction, *N*_0_ *\* f*, (Supplementary Figure 4). We decided to further explore the effects of the latter, as this variable is readily tunable experimentally through changing the number of cells in the reaction or through modulating the expression levels of the nanobody. An analysis that investigated the change in the analyte concentration at which maximum agglutination levels are observed, demonstrated that perturbing these two parameters individually by 10-fold with respect to their nominal values results in the biggest change of all parameter perturbations possible (Figure 2B). In practice, this means that these variables can be used to alter the concentration range of the zone of agglutination. Figure 2C shows the results of these simulations for both variables in their ten-fold perturbation scenarios.

**Figure 2:**
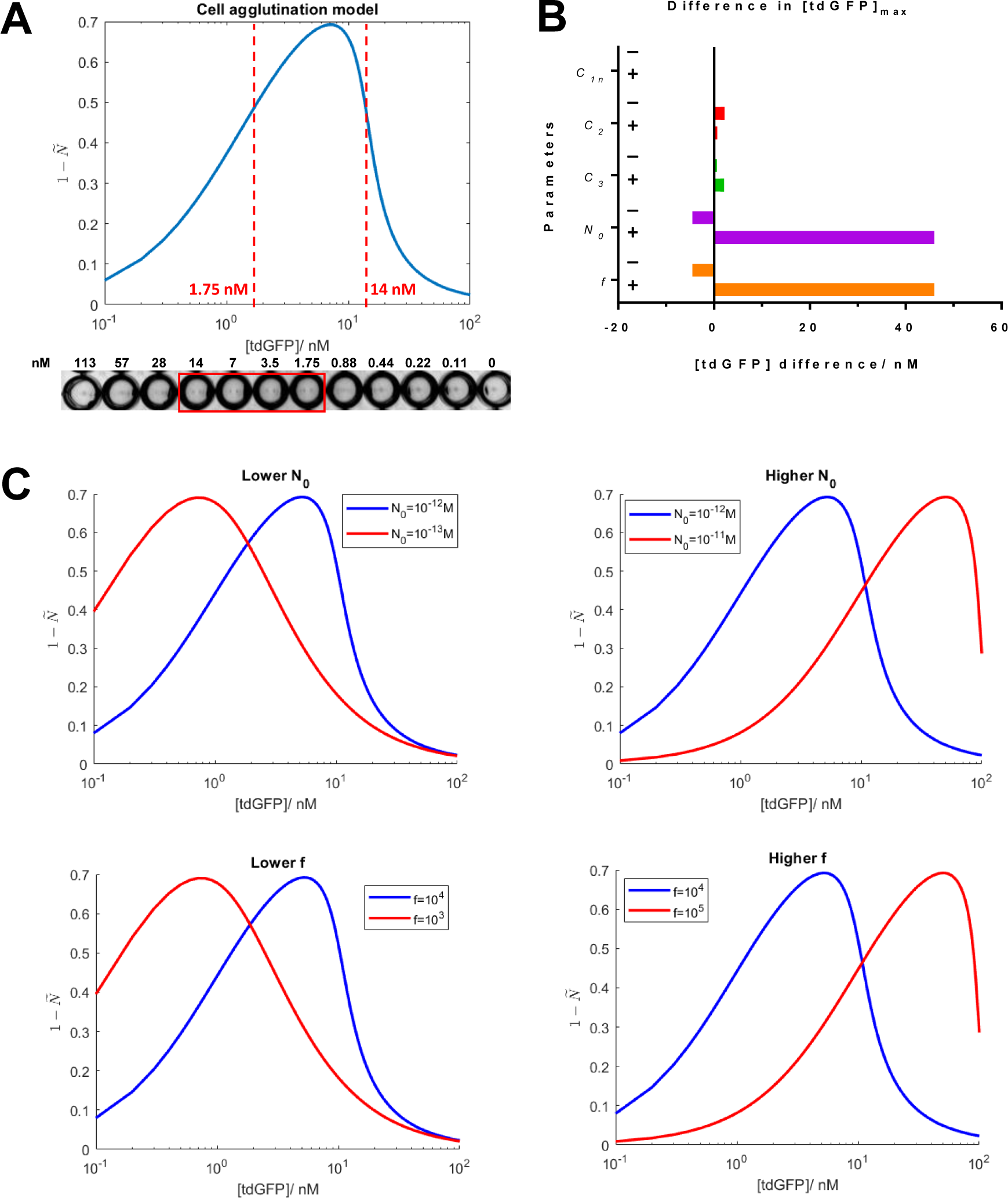
Mathematical modeling and sensitivity analysis of the bacterial agglutination reaction. **A** Computational simulation of the bacterial agglutination reaction for tdGFP analyte concentrations. Y-axis represents the endpoint normalized cell agglutination value 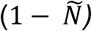 for each analyte concentration when the system is solved numerically over a timespan of 18 hours. The red dotted lines indicate the maximal and minimal tdGFP concentrations (1.75 nM to 14 nM) where cell agglutination was observed in an experimental assay (image of the plate assay at the bottom of the graph, where agglutination-positive samples are indicated with a red rectangle). **B** Bar chart of the change in the concentration of tdGFP analyte at which maximum normalised cell agglutination is observed after parameter perturbation from their nominal values to tenfold higher(+) or lower(-). The parameters are (from top to bottom): normalised agglutination rate constant (*C*1*n*), nanobody binding rate constant (*C*_2_), nanobody dissociation rate constant (*C*_3_), initial concentration of cells (*N*_0_) and number of nanobodies per cell (*f*). **C** [Top graphs] Simulations of the agglutination reaction at either tenfold-lower or tenfold-higher concentrations of bacterial cells (red curve) with respect to the base model (blue curve). [Bottom graphs] Simulations of the agglutination reaction at either tenfold-lower or tenfold-higher levels of nanobody per cell (red curve) with respect to the base model (blue curve).

The total number of nanobodies in the agglutination reaction can be manipulated experimentally by either changing the number of cells in the assay or the number of nanobodies displayed on the surface of each cell. Although changing the number of cells in the reaction is the easier of the two approaches, in practice this is hindered by a lower limit on the concentration of cells that can be detected visually (OD_600nm_ ∼ 0.25, Supplementary Figure 3). While adjusting the number of nanobody per cell is more technically challenging, to facilitate the use of this technology in an easy plug-and-play manner, we developed a library of vectors that allows for different numbers of nanobody displayed on the surface of each cell. For this library, we used three promoters of low, medium and high strengths from the Anderson collection of constitutive promoters (Registry_of_Standard_Biological_Parts 2016), and three ribosome binding site (RBS) designs of low, medium and high strengths designed using the RBS calculator (Salis et al. 2009), leading to nine device variants with different promoter-RBS combinations that were expected to result in different levels of displayed anti-GFP nanobodies.

Initially, we used flow cytometry to measure the levels of nanobody displayed per cell using live cell labeling with monomeric GFP. The data showed a large variation in mean cell fluorescence among device variants (Figure 3, bar charts), indicating varied expression of anti-GFP nanobodies on the surface of the cells. Among the variants, device v02 showed the lowest levels of nanobody display and device v07 showed the highest levels of nanobody display with the rest of the variants showing levels between these extremes. The subsequent use of different variants in the plate agglutination assays with the tdGFP analyte showed shifts in the zone of agglutination that strongly correlate with the nanobody expression levels, confirming the predictions of the simulations using the mathematical model (Figure 3, microplate images). In particular, the dynamic range of device v02 shifted towards lower analyte concentrations allowing the detection of tdGFP analyte at concentrations as low as 450 pM, compared to the previous detection limit of 1.8 nM in the initial prototype.

**Figure 3:**
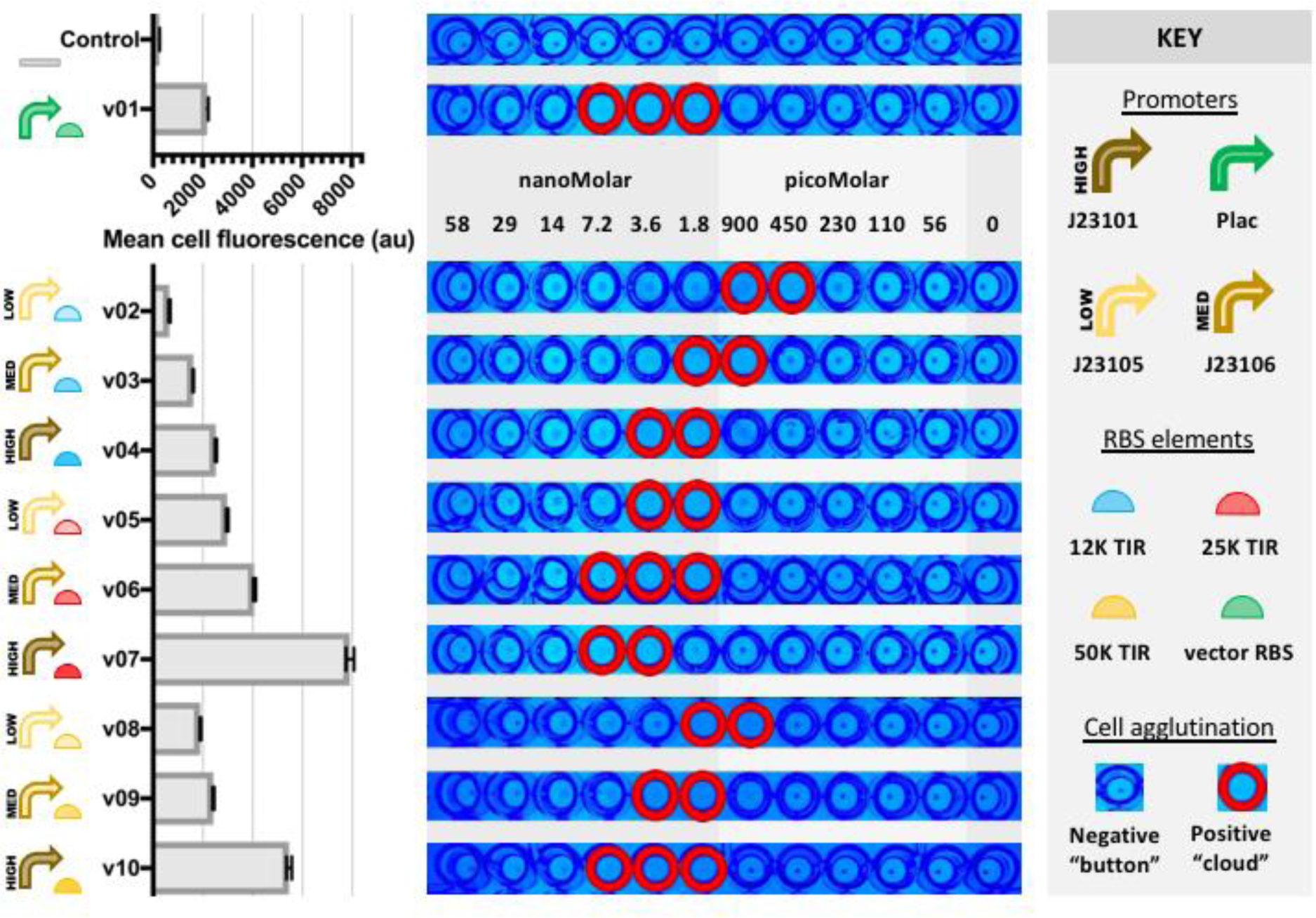
Modulation of the analytical characteristics of the agglutination assay by the control of the nanobody expression levels. The prototype (v01) *E. coli* cells, which have been engineered to display an anti-GFP nanobody on their surface, were genetically modified to produce variants (v02-v10) that express different levels of anti-GFP nanobody. Genetic elements used to control the expression levels of the nanobody display vector are shown next to each variant. The bar chart shows the mean cell fluorescence of each cell variant population when labeled with saturating amounts of purified GFP. Cell fluorescence was recorded by flow cytometry, and error bars show standard deviation for three technical replicates. Images of cell agglutination assay in a microplate, with engineered *E. coli* cells displaying anti-GFP nanobody on their surface after incubation with various tdGFP concentrations. The microplate was back-illuminated with blue light. Control cells did not express any nanobody.

### Bioassay development towards use as a real-world diagnostic test

Human Fibrinogen (hFib) is a glycoprotein with normal blood plasma levels between 1.5 and 4.5 mg/mL. hFib plays a role in blood coagulation when it is converted into fibrin by thrombin protease (Lowe et al. 2004). Elevated levels of fibrinogen in plasma samples are commonly used as a biomarker for determining the risk of cardiovascular disorders in humans (Willis et al. 2004), whereas low levels of fibrinogen in plasma are associated with blood disorders such as disseminated clotting (Lowe et al. 2004). hFib is a heterohexamer, consisting of two dimers of three different polypeptide chains and therefore it is inherently a multivalent antigen.Therefore, we decided to develop a diagnostic test for hFib as a first real-world use case of our new bacterial LAT platform.

We began by constructing two cell lines that surface-display two different anti-hFib nanobody constructs (NbFib1 and NbFib2) with different epitope recognition sites (Salema et al. 2016a). Each nanobody was expressed on the cell surface using the same display vector, and anti-hFib nanobody display was confirmed by flow cytometry and fluorescent microscopy analyses (Supplementary Figure 5). Additionally, we used our mathematical model with updated parameter values specific to the hFib system to predict the range of analyte concentrations at which the agglutination zone would be observed (Figure 4A).

**Figure 4:**
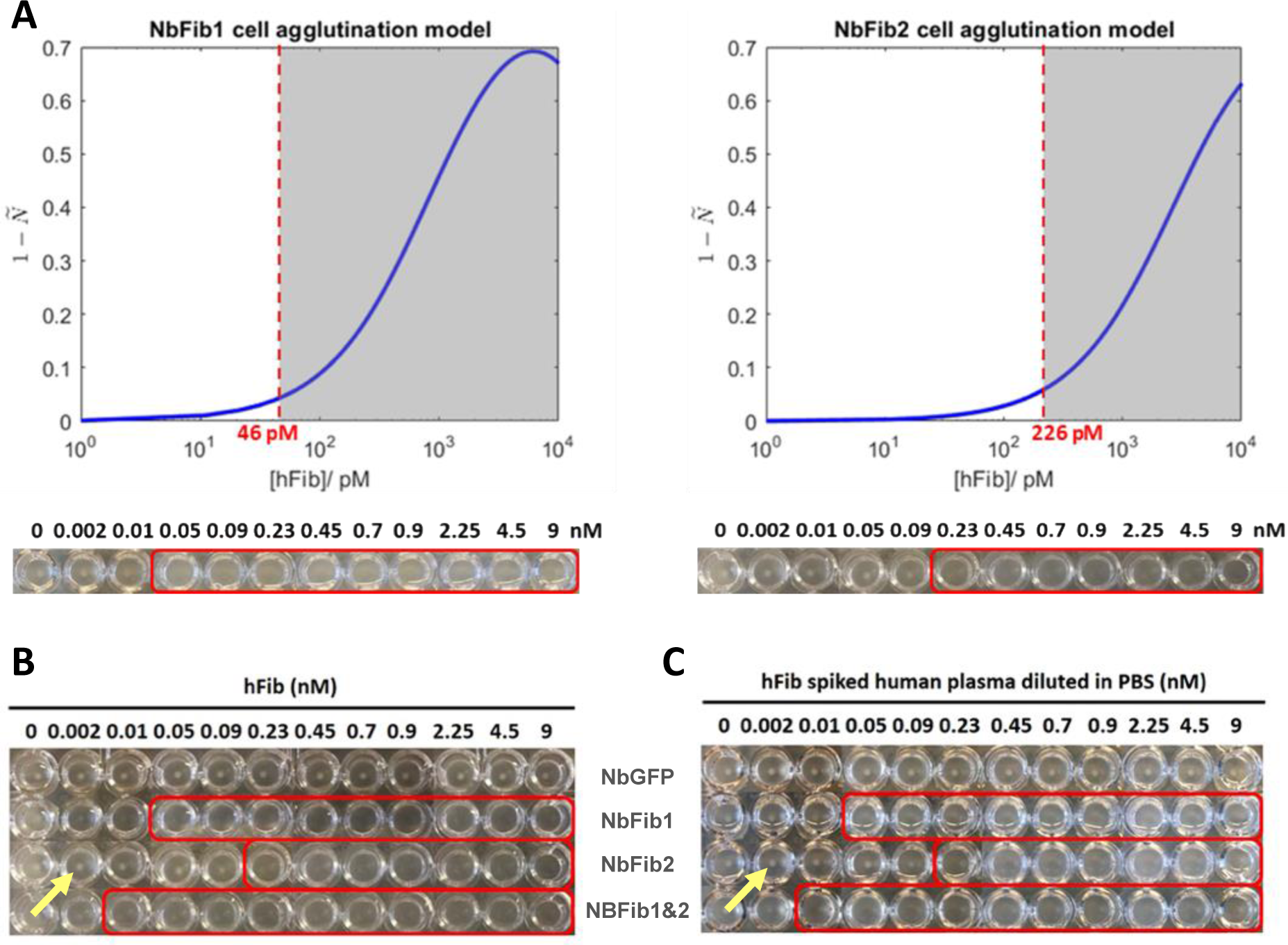
Validating the agglutination assay using Human Fibrinogen (hFib). **A** Computational simulations of the bacterial agglutination reaction for fibrinogen detection for two different anti-fibrinogen biosensor cell strains. Dark region indicates the zone of agglutination and red dotted line indicate the minimal fibrinogen concentration where cell agglutination was observed in experimental assays (image of the plate assay at the bottom of the graphs, where agglutination positive samples are indicated with a red rectangle). **B** Plate agglutination assay of *E. coli* cells expressing anti-hFib nanobodies (NbFib1/2/3). Plate wells included single lines of biosensor cells (plate rows 1-4) or mixed populations of cell lines (rows 5-6) incubated at various hFib concentrations (right). **C** Bacterial agglutination assay of human plasma samples spiked with hFib analyte concentrations. Cells expressing anti-GFP nanobody were used as a negative control. Red boxes indicate zones of bacterial agglutination. Yellow arrows indicate cell pellet formation.

We then verified the designed diagnostic test with purified samples of hFib protein in a microplate agglutination assay format. For each of the two cell lines with nanobodies specific against different hFib epitopes, the assay produced bacterial agglutination zones at the predicted analyte concentration ranges. The detection limit was the lowest for the cell line displaying the NbFib1 nanobody at ∼50 pM hFib in the reaction; the detection limit for the assay using the NbFib2 nanobody was ∼230 pM (Figure 4B). We speculate that these shifts can be attributed to differences in nanobody expression (Supplementary Figure 5). Importantly, a control reaction using cells that display anti-GFP nanobodies did not exhibit any agglutination, providing further evidence that specific interactions between analyte and nanobody elements drive the crosslinking effect.

The availability of nanobodies that bind to distinct epitopes in hFib provided the opportunity to test whether the limit of detection could be further decreased by increasing the propensity for crosslinking through the availability of multiple binding sites (Figure 4B). Indeed, a mixture of the cells expressing NbFib1 and NbFib2, the two that individually had the lowest limit of detection, resulted in a further decrease in the limit of detection to 10 pM hFib.

Finally, to test the robustness of the assay to potential sample matrix effects, we compared the assay results using samples of hFib reconstituted in fibrinogen-deficient human plasma with those reconstituted in phosphate-buffered saline (PBS). As shown in Figure 4C, there was no observable difference between samples prepared in PBS or human plasma samples, suggesting that the assay is robust to the sample background and can be used with real patient samples without interference.

## CONCLUSIONS

In this paper, we describe the development of a biological equivalent to the LAT, an assay widely used in medical diagnostics. Our bioassay includes cells that are genetically engineered to display nanobodies on their surface, analogous to the latex particles coated with antibodies in LATs. The presence of the analyte induces bacterial cross-linking, which is visible to the naked eye and enables facile detection of the target analyte in a manner similar to the readout of the LAT. This expands the repertoire of the types of analytes that can be sensed using whole-cell biosensors to include large molecules such as protein biomarkers or disease agents, which cannot easily be transported into the cell.

Whole-cell biosensors have become an alternative promising tool for the detection of analytes in diverse applications, as they are easy to manipulate, simple and low-cost. Our platform adds a new tool to this arsenal for the extracellular detection of protein analytes. Additionally, our platform is unique in enabling analyte detection without requiring an active cell metabolism. This contrasts with most previous solutions where the output signal is dependent on active cellular metabolism and/or gene expression for reporter protein production (Courbet et al. 2015; Daringer et al. 2014; Ostrov et al. 2017). Lastly, the platform can potentially offer large cost savings to assay manufacturers as the cells autonomously express and surface display nanobodies which removes the need for the costly steps of antibody production, purification and attachment to latex particles surface.

The design of our biological LAT is inherently modular and is readily extensible to the detection of other analytes of interest by exchanging the nanobody that is surface displayed through simple molecular cloning techniques. There are numerous developed nanobodies described in the literature (Harmsen and De Haard 2007). Alternatively, phage-display libraries can be used to select new nanobodies for different targets or to enhance specificity (Tanha et al. 2002; Verheesen et al. 2006)or immunization, e.g. of camels, llamas, or sharks, can be used to isolate new nanobodies if desired (van der Linden et al. 2000). Thus, the technology described here could be used as a platform for the development of a range of assays for different target analytes. For inherently multivalent targets, a nanobody that binds to a single epitope is all that is necessary to develop a new assay. Monomeric targets can be detected by mixing two cell populations expressing nanobodies that bind to different epitopes. Additionally, we have shown that the dynamic range of this diagnostic platform can be easily tuned for the requirement of the particular application, and very low limits of detection are easily achievable. In particular, the limit of detection of our biological LAT assay for hFib was over 29,000-fold lower than that of a piezoelectric agglutination sensor (Chen et al. 2010) and about 13-fold lower than that of an amperometric immunosensor (Campuzano et al. 2014). Using our mathematical model in combination with the developed library of variants of nanobody expression, new diagnostic tests can be conveniently designed for target analyte concentrations at will.

An important consideration for the adoption of new technologies is ease-of-use and seamless integration into existing professional workflows. Our biological LAT platform is underscored by the simplicity of its design and gives a visual readout that is easy to interpret. In addition, it fits within existing clinical diagnostic frameworks since the LAT is used in clinical laboratories to detect a number of pathogens including *H. influenzae* (Heikkilä et al. 1987), *N. meningitides* (Leinonen and Herva 1977), *P.* and *S. pneumoniae* (Smith and Washington 1984) and therefore, it could readily be adopted by existing personnel with minimal training required to implement it.

## ACKNOWLEDGMENTS

We would like to thank the following funding sources: EPSRC EP/K038648/1 (KP, PF); BBSRC CASE Studentship (NK); Royal Thai Government Scholarship (PR), MICIIU BIO2017-89081R (LAF); EPSRC EP/P009352/1 & EP/M002187/1(GBS); Imperial College President’s PhD Scholarship (HEL).

## AUTHORS CONTRIBUTIONS

Study design: NK, KP, PF; Experimental execution and data analysis: NK, PR, HEL; Mathematical modeling: HEL, GBS; Nanobody development: VS, LAF; Writing manuscript: NK, KP in consultation with the other authors.

## REFERENCES

Assicot, M., Gendrel, D., Carsin, H., Raymond, J., Guilbaud, J., Bohuon, C., 1993. High serum procalcitonin concentrations in patients with sepsis and infection. Lancet (London, England) 341(8844), 515–518.

Borrebaeck, C.A., 2000. Antibodies in diagnostics - from immunoassays to protein chips. Immunology today 21(8), 379–382.

Campuzano, S., Salema, V., Moreno-Guzman, M., Gamella, M., Yanez-Sedeno, P., Fernandez, L.A., Pingarron, J.M., 2014. Disposable amperometric magnetoimmunosensors using nanobodies as biorecognition element. Determination of fibrinogen in plasma. Biosensors & bioelectronics 52, 255–260.

Cavallari, M., 2017. Rapid and Direct VHH and Target Identification by Staphylococcal Surface Display Libraries. International journal of molecular sciences 18(7).

Chen, Q., Hua, X., Fu, W., Liu, D., Chen, M., Cai, G., 2010. Quantitative determination of fibrinogen of patients with coronary heart diseases through piezoelectric agglutination sensor. Sensors 10(3), 2107–2118.

Courbet, A., Endy, D., Renard, E., Molina, F., Bonnet, J., 2015. Detection of pathological biomarkers in human clinical samples via amplifying genetic switches and logic gates. Science translational medicine 7(289), 289ra283.

Daar, A.S., Thorsteinsdottir, H., Martin, D.K., Smith, A.C., Nast, S., Singer, P.A., 2002. Top ten biotechnologies for improving health in developing countries. Nature genetics 32(2), 229–232.

Daringer, N.M., Dudek, R.M., Schwarz, K.A., Leonard, J.N., 2014. Modular extracellular sensor architecture for engineering mammalian cell-based devices. ACS synthetic biology 3(12), 892–902.

Daugherty, P.S., Chen, G., Olsen, M.J., Iverson, B.L., Georgiou, G., 1998. Antibody affinity maturation using bacterial surface display. Protein engineering 11(9), 825–832.

Dolgosheina, E.B., Karulin, A., Bobylev, A.V., 1992. A kinetic model of the agglutination process. Mathematical biosciences 109(1), 1–10.

Fleetwood, F., Devoogdt, N., Pellis, M., Wernery, U., Muyldermans, S., Stahl, S., Lofblom, J., 2013. Surface display of a single-domain antibody library on Gram-positive bacteria. Cellular and molecular life sciences: CMLS 70(6), 1081–1093.

Francisco, J.A., Campbell, R., Iverson, B.L., Georgiou, G., 1993. Production and fluorescence-activated cell sorting of Escherichia coli expressing a functional antibody fragment on the external surface. Proc Natl Acad Sci U S A 90(22), 10444–10448.

Fuchs, P., Breitling, F., Dubel, S., Seehaus, T., Little, M., 1991. Targeting recombinant antibodies to the surface of Escherichia coli: fusion to a peptidoglycan associated lipoprotein. Bio/technology (Nature Publishing Company) 9(12), 1369–1372.

Garcia-Molla, V., 2017a. Sensitivity Analysis for ODEs and DAEs. p. Solves ODE/DAE systems (as ODE15s solver) and studies dependence of solutions wrt parameters., version 1.2.0.0 ed. Matlab, File Exchange - MATLAB Central.

Garcia-Molla, V., 2017b. Sensitivity Analysis for ODEs and DAEs. p. Sensitivity Analysis for ODEs and DAEs, version 1.2.0.0 ed. Matlab, File Exchange - MATLAB Central.

Giljohann, D.A., Mirkin, C.A., 2009. Drivers of biodiagnostic development. Nature 462(7272), 461–464.

Goers, L., Ainsworth, C., Goey, C.H., Kontoravdi, C., Freemont, P.S., Polizzi, K.M., 2017. Whole-cell Escherichia coli lactate biosensor for monitoring mammalian cell cultures during biopharmaceutical production. Biotechnology and bioengineering 114(6), 1290–1300.

Goers, L., Kylilis, N., Tomazou, M., Wen, K., Freemont, P., Polizzi, K., 2013. Engineering Microbial Biosensors. In: Colin, H., Anil, W. (Eds.), Microbial Synthetic Biology, pp. 119–156, 1st ed. Elsevier.

Harmsen, M.M., De Haard, H.J., 2007. Properties, production, and applications of camelid single-domain antibody fragments. Applied Microbiology and Biotechnology 77(1), 13–22.

Haverkate, F., Thompson, S.G., Pyke, S.D., Gallimore, J.R., Pepys, M.B., 1997. Production of C-reactive protein and risk of coronary events in stable and unstable angina. European Concerted Action on Thrombosis and Disabilities Angina Pectoris Study Group. Lancet (London, England) 349(9050), 462–466.

Heikkilä, R., Takala, A., Käyhty, H., Leinonen, M., 1987. Latex agglutination test for screening of Haemophilus influenzae type b carriers. J Clin Microbiol 25(6), 1131–1133.

Horswell, J., Dickson, S., 2003. Use of biosensors to screen urine samples for potentially toxic chemicals. Journal of analytical toxicology 27(6), 372–376.

Leinonen, M., Herva, E., 1977. The latex agglutination test for the diagnosis of meningococcal and haemophilus influenzae meningitis. Scandinavian journal of infectious diseases 9(3), 187– 191.

Looger, L.L., Dwyer, M.A., Smith, J.J., Hellinga, H.W., 2003. Computational design of receptor and sensor proteins with novel functions. Nature 423(6936), 185–190.

Lowe, G.D., Rumley, A., Mackie, I.J., 2004. Plasma fibrinogen. Annals of clinical biochemistry 41(Pt 6), 430–440.

Muyldermans, S., 2013. Nanobodies: natural single-domain antibodies. Annual review of biochemistry 82, 775–797.

Network, W.G.I.S., 2011. Manual for the laboratory diagnosis and virological surveillance of influenza. World Health Organization.

NICE, 2014. Pneumonia in adults: diagnosis and management recommendations | Guidance and guidelines | NICE. 1.1 Presentation with lower respiratory track infection. National Institute for Health and Care Excellence.

Ohman, E.M., Armstrong, P.W., Christenson, R.H., Granger, C.B., Katus, H.A., Hamm, C.W., O’Hanesian, M.A., Wagner, G.S., Kleiman, N.S., Harrell, F.E., Jr., Califf, R.M., Topol, E.J., 1996. Cardiac troponin T levels for risk stratification in acute myocardial ischemia. GUSTO IIA Investigators. The New England journal of medicine 335(18), 1333–1341.

Ortega-Vinuesa, J.L., Bastos-Gonzalez, D., 2001. A review of factors affecting the performances of latex agglutination tests. Journal of biomaterials science. Polymer edition 12(4), 379–408.

Ostrov, N., Jimenez, M., Billerbeck, S., Brisbois, J., Matragrano, J., Ager, A., Cornish, V.W., 2017. A modular yeast biosensor for low-cost point-of-care pathogen detection. Science advances 3(6), e1603221.

Price, C.P., 2001. Agglutination Techniques for Detecting Antigen–Antibody Reactions. eLS. John Wiley & Sons, Ltd.

Registry_of_Standard_Biological_Parts, 2016. Promoters/Catalog/Anderson-parts.igem.org.

Roda, A., Pasini, P., Mirasoli, M., Guardigli, M., Russo, C., Musiani, M., Baraldini, M., 2001. Sensitive determination of urinary mercury(II) by a bioluminescent transgenic bacteria-based biosensor. ANAL LETT 34(1), 29–41.

Salema, V., Fernandez, L.A., 2017. Escherichia coli surface display for the selection of nanobodies. Microbial biotechnology.

Salema, V., Lopez-Guajardo, A., Gutierrez, C., Mencia, M., Fernandez, L.A., 2016a. Characterization of nanobodies binding human fibrinogen selected by E. coli display. Journal of biotechnology.

Salema, V., Manas, C., Cerdan, L., Pinero-Lambea, C., Marin, E., Roovers, R.C., van Bergen En Henegouwen, P.M., Fernandez, L.A., 2016b. High affinity nanobodies against human epidermal growth factor receptor selected on cells by E. coli display. mAbs, 0.

Salema, V., Marin, E., Martinez-Arteaga, R., Ruano-Gallego, D., Fraile, S., Margolles, Y., Teira, X., Gutierrez, C., Bodelon, G., Fernandez, L.A., 2013. Selection of single domain antibodies from immune libraries displayed on the surface of E. coli cells with two beta-domains of opposite topologies. PLoS one 8(9), e75126.

Salis, H.M., Mirsky, E.A., Voigt, C.A., 2009. Automated Design of Synthetic Ribosome Binding Sites to Precisely Control Protein Expression. Nat Biotechnol 27(10), 946–950.

Smith, S.K., Washington, J.A., 1984. Evaluation of the Pneumoslide latex agglutination test for identification of Streptococcus pneumoniae. J Clin Microbiol 20(3), 592–593.

Tanha, J., Dubuc, G., Hirama, T., Narang, S.A., MacKenzie, C.R., 2002. Selection by phage display of llama conventional V(H) fragments with heavy chain antibody V(H)H properties. Journal of immunological methods 263(1-2), 97–109.

Thompson, I.M., Pauler, D.K., Goodman, P.J., Tangen, C.M., Lucia, M.S., Parnes, H.L., Minasian, L.M., Ford, L.G., Lippman, S.M., Crawford, E.D., Crowley, J.J., Coltman, C.A., Jr., 2004. Prevalence of prostate cancer among men with a prostate-specific antigen level < or =4.0 ng per milliliter. The New England journal of medicine 350(22), 2239–2246.

Turner, K., Xu, S., Pasini, P., Deo, S., Bachas, L., Daunert, S., 2007. Hydroxylated polychlorinated biphenyl detection based on a genetically engineered bioluminescent whole-cell sensing system. Analytical chemistry 79(15), 5740–5745.

Urdea, M., Penny, L.A., Olmsted, S.S., Giovanni, M.Y., Kaspar, P., Shepherd, A., Wilson, P., Dahl, C.A., Buchsbaum, S., Moeller, G., Hay Burgess, D.C., 2006. Requirements for high impact diagnostics in the developing world. Nature 444 Suppl 1, 73–79.

van der Linden, R., de Geus, B., Stok, W., Bos, W., van Wassenaar, D., Verrips, T., Frenken, L., 2000. Induction of immune responses and molecular cloning of the heavy chain antibody repertoire of Lama glama. Journal of immunological methods 240(1-2), 185–195.

van der Meer, J.R., Belkin, S., 2010. Where microbiology meets microengineering: design and applications of reporter bacteria. Nature reviews. Microbiology 8(7), 511–522.

Verheesen, P., Roussis, A., de Haard, H.J., Groot, A.J., Stam, J.C., den Dunnen, J.T., Frants, R.R., Verkleij, A.J., Theo Verrips, C., van der Maarel, S.M., 2006. Reliable and controllable antibody fragment selections from Camelid non-immune libraries for target validation. Biochimica et biophysica acta 1764(8), 1307–1319.

Webb, A.J., Kelwick, R., Doenhoff, M.J., Kylilis, N., MacDonald, J.T., Wen, K.Y., McKeown, C., Baldwin, G., Ellis, T., Jensen, K., Freemont, P.S., 2016. A protease-based biosensor for the detection of schistosome cercariae. Scientific reports 6, 24725.

Willis, A., Jung, E.J., Wakefield, T., Chen, X., 2004. Mutant p53 exerts a dominant negative effect by preventing wild-type p53 from binding to the promoter of its target genes. Oncogene 23(13), 2330–2338.

